# Vestibular mapping of the naturalistic head-centered motion spectrum

**DOI:** 10.1101/2023.06.12.544528

**Authors:** Matthias Ertl, Peter zu Eulenburg, Marie Woller, Ümit Mayadali, Rainer Boegle, Marianne Dieterich

**Affiliations:** Department of Psychology, University of Bern, Switzerland; Department of Neurology, Ludwig-Maximilians-Universität München, Germany; German Center for Vertigo and Balance Disorders (IFBLMU), Ludwig-Maximilians-Universität München, Germany; Graduate School of Systemic Neuroscience, Ludwig-Maximilians-Universität München, Germany; Munich Cluster for Neurology (SyNergy), Munich, Germany

**Author notes:** Corresponding Author: Dr. rer. biol. hum. Matthias Ertl University of Bern, Switzerland Department of Psychology Fabrikstrasse 8 3012 Bern.

**Keywords:** cingulate sulcus visual, direction dependency, vestibular anisotropy, vestibular evoked potentials, passive motion

## Abstract

**BACKGROUND:** Naturalistic head accelerations can be used to elicit vestibular evoked potentials (VestEPs). These potentials allow for analysis of cortical vestibular processing and its multi-sensory integration with a high temporal resolution.

**METHODS:** We report the results of two experiments in which we compared the differential VestEPs elicited by randomized translations, rotations, and tilts in healthy subjects on a motion platform.

**RESULTS:** An event-related potential (ERP) analysis revealed that established VestEPs were verifiable in all three acceleration domains (translations, rotations, tilts). A further analysis of the VestEPs showed a significant correlation between rotation axes (yaw, pitch, roll) and the amplitude of the evoked potentials. We found increased amplitudes for rotations in the roll compared to the pitch and yaw plane. A distributed source localization analysis showed that the activity in the cingulate sulcus visual (CSv) area best explained direction-dependent amplitude modulations of the VestEPs, but that the same cortical network (posterior insular cortex, CSv) is involved in processing vestibular information, regardless of the motion direction.

**CONCLUSION:** The results provide evidence for an anisotropic, direction-dependent processing of vestibular input by cortical structures. The data also suggest that area CSv plays an integral role in ego-motion perception and interpretation of spatial features such as acceleration direction and intensity.

## Introduction

Despite its everyday importance for locomotion and navigation, the cortical processing and integration of vestibular input in humans is still not as well understood as all other sensory systems. Most of our knowledge of vestibular cortical processing has been achieved by non-invasive functional magnetic resonance imaging (fMRI) and positron emission tomography (PET) studies [3,4,9,32,37,47,52]. From these neuroimaging studies it is known that vestibular input is processed by a network of distinct brain areas including the parieto-opercular region, the sylvian fissure, the superior temporal gyrus, and the cingulate cortex [34,59]. These regions were found to be vestibular core regions in primates as well as non-primate species [33].

Behavioral and psychophysical experiments demonstrated that the human vestibular system samples and represents the three-dimensional (3D) head-centered space in an anisotropic fashion. For example, the vestibular thresholds for translations vary with respect to the acceleration axis and are increased for sideways compared to fore/aft accelerations in healthy subjects [7]. Additionally, systematic biases in human heading estimation have been reported in naturalistic acceleration experiments. These direction-specific modulations of the vestibular system might contribute to anisotropies observed in tasks that involve higher vestibular functions, such as the spatial representation of big 3D objects [6,13] and 3D navigation [5,60], which are also biased with respect to specific directions.

Complementary to neuroimaging experiments using PET and MRI, a few EEG studies on the processing of vestibular input in humans have been performed over the last decades [11,14,18,28,55]. These studies tried to exploit the high temporal resolution of the EEG to investigate the neural cortical correlates of vestibular stimulation. Studies using rotatory chairs showed that a series of short (< 10 ms), middle (10 – 30 ms), and long (> 30 ms) latency vestibular evoked potentials (VestEPs) can be detected on the scalp during and after whole-body accelerations around the vertical axis [2,11,23,30]. Other stimulation approaches, such as loud click-sounds [27,35,55], mechanic impulses [53,54], or vestibular nerve stimulation during surgical interventions [57], were also successfully used to elicited VestEPs with short, middle, and long latencies. However, the applied stimuli and analysis were quite heterogeneous across the studies and many of the reported findings contradict each other. In particular, the described late VestEPs and topographies vary widely across the studies [15,46,50].

In addition to event-related potentials, a second supplementary branch of EEG analysis has evolved over the last decades. Instead of focusing only on phenomena in the time domain, the attention of many theories and studies shifted toward the description of oscillation patterns in the frequency or time-frequency domain. Consequently, many brain (mal-) functions have been linked to specific frequency bands [21]. Time-frequency analyses have also already been performed in the context of vestibular processing. A pioneering study reported a significant power increase in the delta-band and a power decrease in the theta-and alpha-band during stimulation of the horizontal canal [2]. By applying a rotation around the vertical axis for multiple seconds a more recent study [18] found an alpha-band suppression at the vertex and a beta-band suppression over medial fronto-parietal scalp regions. Both studies were conducted on rotatory chairs using sinusoidal motion profiles. However, reports of increased beta-band activity elicited by impulse-like translational accelerations can be found in the literature [15].

Driven by the heterogeneous and partly contradictory results on VestEPs and oscillation patterns reported by previous studies, we conducted two experiments to systematically investigate the cortical responses evoked by rotations, translations, and pitch/roll tilts. Based on animal research [1,31], we did not expect differences in the cortical representation of the time-courses of translations and tilts. However, due to previous findings regarding an anisotropic vestibular perception and representation of 3D space [60] in humans, we expected to find significant differences between acceleration trajectories during the cortical processing for different directions.

## Methods

Fifty healthy subjects (HS) were included in the study; five participated in both recordings while all other subjects differed between the two experiments. The first experiment focused on the effects of tilts versus translations. The second experiment aimed to investigate the effects of rotations. The acceleration profiles in our experiments and the analyses were kept as similar as possible to the recently published study on VestEPs during translational accelerations [14,15]. All HS had no prior history of neurological or neuro-otological disease and did not take any medication on a regular basis. HS gave their informed written consent and were paid for participation. The local ethics committee approved the study.

### Experiment 1 – Tilt and Translation

#### Subjects

For this experiment, 25 HS were measured. Three participants did not complete the experiment due to motion sickness; another four datasets had to be excluded based on visual inspection of the raw data from the analysis due to insufficient overall data quality. Recording EEG in an electrically noisy environment (not a shielded room) during passive movements is a challenging task. The accelerations can cause severe artifacts, particularly if the headphones or neck brace are not positioned carefully enough. The excluded datasets suffered from major artifacts (signal loss at electrodes, huge overall noise, and selection of wrong recording parameters) for large portions of the recorded data. The remaining cohort (n=18; 12 females) had a mean age of 30.6 years (SD = 6.9). Two of the HS were left-handed and 16 right-handed according to the 10-item inventory of the *Edinburgh test* [38].

#### Stimuli and Procedure

Subjects were seated with the head-vertical axis aligned with the gravitation vector with the heads slightly tilted downwards and restrained in a padded racing car seat with straps mounted on a 6-degree-of-freedom motion platform (Moog© 6DOF2000E). Motion platforms are one of few possibilities for stimulating the vestibular system in a naturalistic way and without introducing a multi-sensory conflict [12]. To mask motion-related sounds generated by the platform, HS wore noise-cancelling headphones and white noise (90 dB SPL) was presented throughout the recordings. To minimize delays between platform motion and consecutive head movement participants wore a neck brace throughout the experiment. Because of its shape, the adjustable neck brace (Stifneck® Select™ Adjustable Collar, Laerdal Medical) prevented the participants from rotating their head relative to the trunk. In this experiment, the platform performed a total of 960 accelerations. The accelerations for the whole session were split into three randomized blocks of 320 stimuli with a 90 s pause between the blocks. Eight different motion profiles (Fig. 1) stimulating the utricles (translation left/right (1), translation fore/aft (2)), the anterior and posterior semicircular canal (pitch tilt (3), roll tilt (4)), a combination of canals and otolith organs (roll tilt plus translation left/right (5), pitch tilt plus translation fore/aft (6)), and a profile aiming to isolate the canals by cancelling the otolith signals (roll tilt minus translation left/right (7), pitch tilt minus translation fore/aft (8)) were used in this experiment. Stimuli 5 – 8 consisted of a combination of a rotation and a translation. The center for the rotation was set at about the height of the ears (1.2 m higher than the default rotation center of the platform). The appropriate combination of a tilt and translation component (roll - translation, pitch - translation) allows for a cancellation of the net acceleration detected by the otolith organs [1,19,31], other combinations (roll + translation, pitch + translation) result in an acceleration detectable by all sub-components of the vestibular end organ. The maximum angle for roll movements was ±5° and ±2.5° for pitch movements and the maximum translational displacement during combined tilt translation movements was 8.33 cm (left/right) and 4.17 cm (fore/aft). Since we were interested in evoked potentials, we used stimuli profiles with well-defined onsets and offsets. The maximum accelerations as measured at the backside of the racing car seat were 0.52 g for left/right and 0.25 g for fore/aft translations, 113 deg/s^2^ for roll-tilt, and 153 deg/s^2^ for the pitch-tilt movements. Similar acceleration profiles were validated in previous studies on VestEPs elicited by translations [14,15]. Samples of the position signals sent to the motion platform and the resulting acceleration profiles for all eight stimuli are visualized in Fig 1. The order of the eight different conditions was randomized and therefore unpredictable for the subject. In addition, the inter-stimulus intervals (ISI) were randomized with an overall mean ISI of 1000 ms ± 250 ms. Two 3D accelerometers (Brain Products Gilching, Germany) recorded the platform motion during the experiment. The accelerometer signals were transmitted to the EEG recorder via an additional amplifier. During the experiment, the subjects had to close their eyes and were blindfolded to ensure absence of visual heading information. Subjects were asked to passively experience the acceleration.

**Fig. 1:**
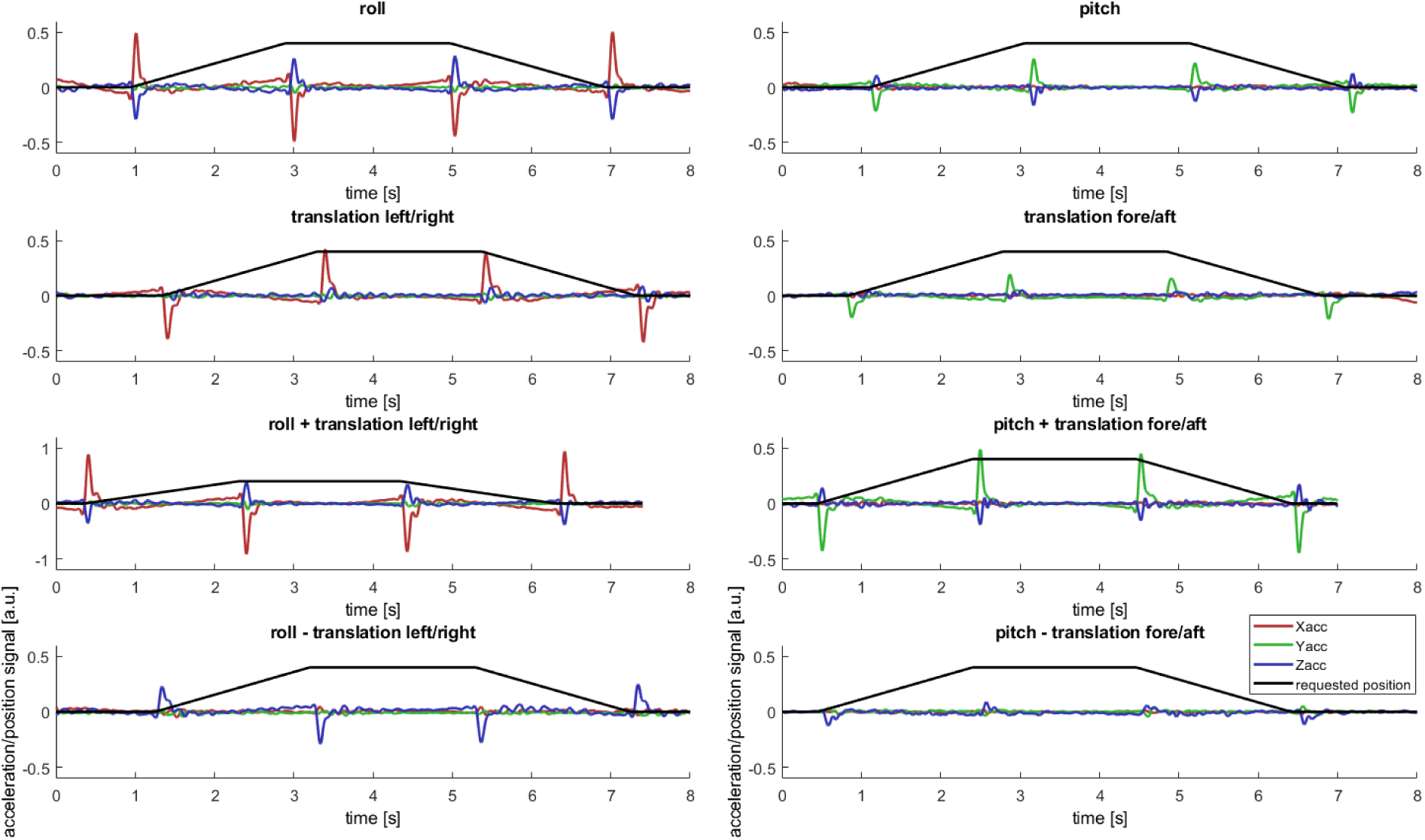
Visualization of sample motion profiles used in experiment 1. The black line represents the position signal sent to the motion platform. The red, green, and blue lines represent the acceleration signals in x, y, z direction as detected by the accelerometer attached to the back of the chair close to the subject’s head. The onset/offset of each motion is characterized by a steep acceleration. Compared to the pure translations (second row), the tilt+translation profiles increase the translational acceleration component (third row), while the tilt-translation profiles cancel as intended the translational acceleration component of the profile (fourth row).

### Experiment 2 – Rotations

#### Subjects

From the dataset of 30 HS one dataset had to be removed from the analyses due to poor data quality. We used the same exclusion criteria as in experiment 1. The larger exclusion rate in experiment 1 is likely caused by the fact that the experiment took longer, which increases the chance for artifacts. The remaining 29 HS (17 female) had a mean age of 25.7 years (SD: 7.21 years). Twenty-five subjects were 100% right-handed, two 100% left-handed. The remaining two subjects were ambidextrous scoring 10% and 70% right-handed on the 10-item inventory [38].

#### Stimuli and Procedure

For the second experiment, the motion platform was programmed to perform 160 rotatory accelerations in each of the three main planes (yaw, pitch, roll). The maximum accelerations experienced by the subjects were 102 deg/s^2^. Samples of the position signal sent to the motion platform and the resulting accelerations as detected by the sensors attached to the platform are visualized in Fig. 2. To minimize motion-related eye-movements, such as the vestibulo-ocular reflex, subjects were asked to continuously fixate a midline near point displayed on a screen about 0.5 m straight ahead. The screen was attached to the platform. The HS were instructed to passively experience the rotations. Since subjects were restrained with their head slightly tilted downwards, the yaw rotations were predominately sensed by the horizontal canals, while the roll and pitch rotations were sensed by both the anterior and posterior canals.

**Fig. 2:**
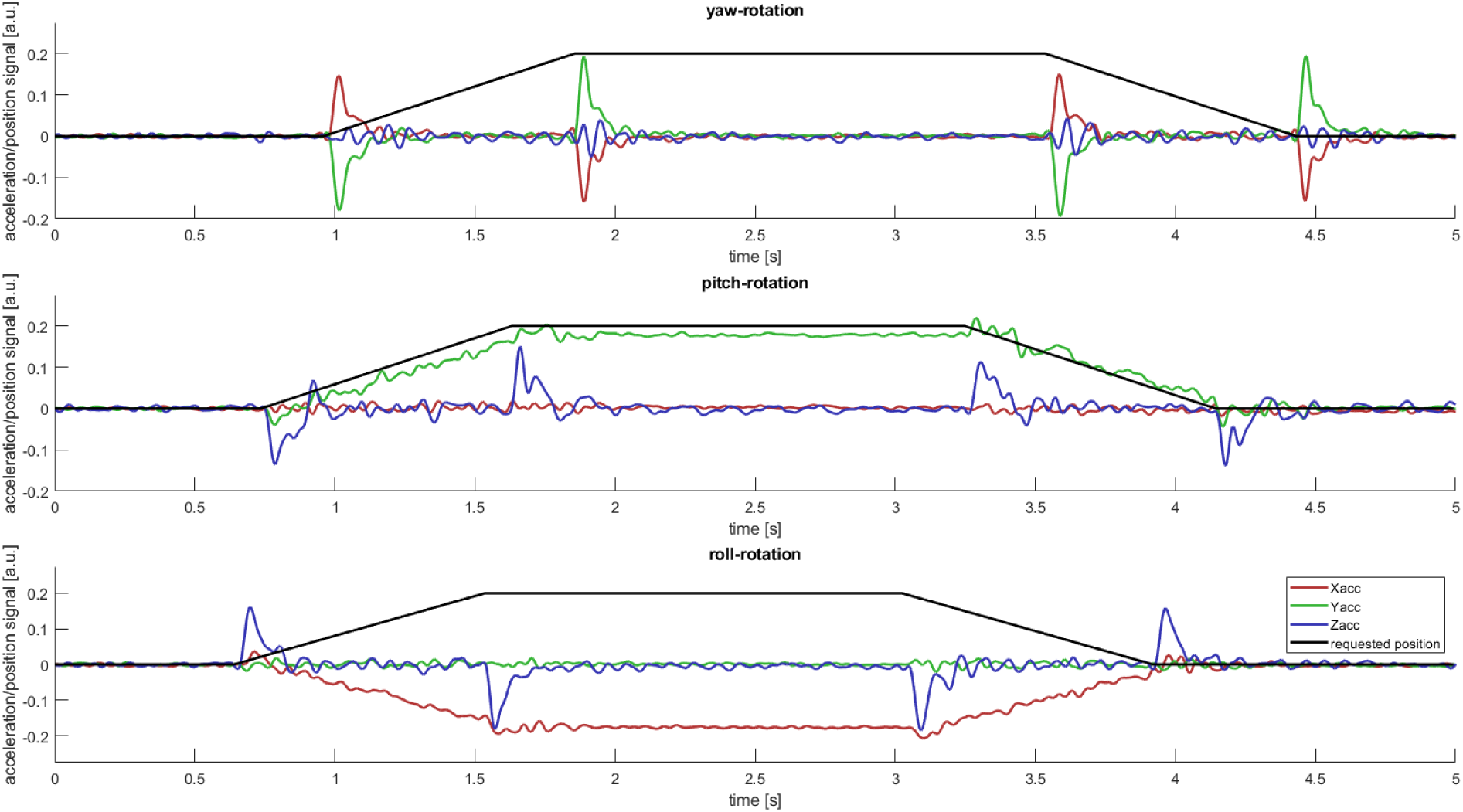
Visualization of sample motion profiles used in experiment 2. The black line represents the position signal sent to the motion platform. The red, green, and blue lines represent the translational components of the acceleration signals in x, y, z direction as detected by the accelerometer attached to the motion platform (∼1.2 m below the rotation center). The onset/offset set of each motion is characterized by a steep acceleration. Yaw-rotations were detected by the two axes of the accelerometer (x, y) positioned in the plane perpendicular to the rotation axis. As the accelerometers only capture the translational components and do not match the angular acceleration sensed by the vestibular organs, the ordinates are shown as arbitrary units [a.u.]. For pitch and roll movements the peaks are accompanied by a drift in one of the axes caused by the position change relative to the gravitational vector of the earth. Therefore, pitch and roll movements also stimulated the otolith organs.

#### EEG recording and preprocessing (both experiments)

The EEG was recorded using 32 active electrodes positioned according to the international 10/20 system with the reference electrode at position FCz. Data were collected at a rate of 1000 samples per second (sps). All impedances were kept below 5 kΩ. The raw EEG data were collected using a 32 EEG channel *BrainAmp MR+* amplifier and an *ExG MR* amplifier for recording data from the accelerometers. All data were recorded using the *Brain Vision Recorder* software Version 1.20 and subsequently analyzed using the *Brain Vision Analyzer* software Version 2.1 (RRID:SCR_002356; Brain Products, Gilching, Germany). Initially the acceleration signals were analyzed using software programmed in-house. The algorithm searched for the maximum accelerations in all conditions and wrote markers to the EEG files. After that, a band-pass filter from 0.1 to 45 Hz was applied to the raw EEG data. After filtering, the data were down sampled to 250 sps and re-referenced to a common average reference. FCz was recovered as regular channel. Artifacts were corrected using an independent component analysis (ICA) approach [20,25]. The independent components were visually inspected, and artifacts were identified using the topographies and the time courses of the components. Visual artifact removal is, despite the development of (semi-)automated procedures [44], a widely used procedure and artifacts can be identified with some experience. Components clearly reflecting artifacts, such as eye-blinks and muscle artifacts were removed [22,25]. A recent study investigated the reliability of ICA as a correction method in a saccade contaminated dataset and found it to be a useful approach [10].

#### ERP

The preprocessed datasets were segmented into epochs from -500 to 800 ms relative to the maximum acceleration. Before averaging, a baseline correction based on the entire pre-stimulus interval was performed. Segments containing artifacts (low activity < .5µV, large amplitude > ±200µV, large amplitude range > 200µV, large amplitude steps > 50µV/µs) were identified by an automatic artifact detection and excluded from the following averaging procedure. Based on the criteria 6.04 % of the trials were excluded in the rotation experiment and 7.40 % in the tilt/translation experiment. For all subjects, separate averages were calculated for every condition. Furthermore, an average containing all trials was calculated and grand averages across all subjects were calculated.

#### Time-Frequency-Analysis

To analyze evoked effects in the frequency domain a wavelet transformation was performed on the averaged time course of every subject. The transformation was applied for the frequency range from 8 to 40 Hz. The pre-stimulus interval (-500 to 0 ms) was used to normalize the output. The frequency range was chosen because our previous study [15] had shown a distinctly evoked response in the beta-band.

#### Source localization

Today EEG source localization is established as a useful approach to reveal sources of evoked potentials and oscillations. One of the most successful algorithms solving the ill-posed problem of source reconstruction is the exact low resolution brain electromagnetic algorithm (*eLORETA; RRID:SCR013830*), a non-linear extension of *sLORETA* [41,42]. *LORETA* is a frequently used algorithm for distributed source localization and has been validated by studies using combined EEG-fMRI [36] and EEG-PET [45] data. *LORETA* has also successfully been used to target activity in deep-lying structures such as the insular cortex [24,29,39,43,48,49,56]. Here we used the *eLORETA* software, which is freely available through the LORETA-website (http://www.uzh.ch/keyinst/loreta.htm). A detailed description of the technical aspects of the algorithms can also be read at the same location.

For this study, the averaged ERPs of every subject were exported to the *LORETA* software package. A transformation matrix was applied to transform every ERP as recorded on the scalp into the source space. To reveal the sources of the VestEPs, a 400 ms baseline ranging from -500 to -100 ms relative to the maximum acceleration was contrasted against the interval in which the VestEPs typically occur (0 – 310 ms).

To identify the sources of the evoked vestibular beta band response (EVBBR) we calculated the time-varying cross-spectrum for every subject and all conditions (translation/rotation/tilt). The spectra were then transformed into source space and a contrast EVBBR-baseline for every motion category (translation, tilt, and rotation) was calculated.

## Results

### ERPs – translations

The averaged time course across all subjects showed three distinct vestibular evoked potentials for translational motion profiles. The first detectable peak had a latency of 31.8 ms (SD = 9.45 ms) and a mean amplitude of 1.07 µV (SD = 0.55 µV). The second component, the N1 reached its mean peak of -6.17 µV (SD = 3.20 µV) 79.8 ms (7.57 ms) after the acceleration maximum. The third component (P2) had a mean amplitude of 5.06 µV (SD = 1.80 µV) and peaked at a latency of 181 ms (SD = 55.7 ms).

### ERPs – tilts

The grand average of VestEPs elicited by tilts around the pitch and roll axes had a time course like those evoked by translations (Fig. 3). The P1 had an amplitude of 0.62 µV (SD = 0.37 µV) and peaked 29.3 ms (SD = 10.8 ms) after the acceleration maximum. The N1 potential reached its maximum amplitude of -3.38 µV (SD = 1.72 µV) with a latency of 83.8 ms (SD = 11.2 ms). The P2 had a mean amplitude of 3.55 µV (SD = 1.41 µV) and peaked with a latency of 203 ms (SD = 49.2 ms).

**Fig. 3:**
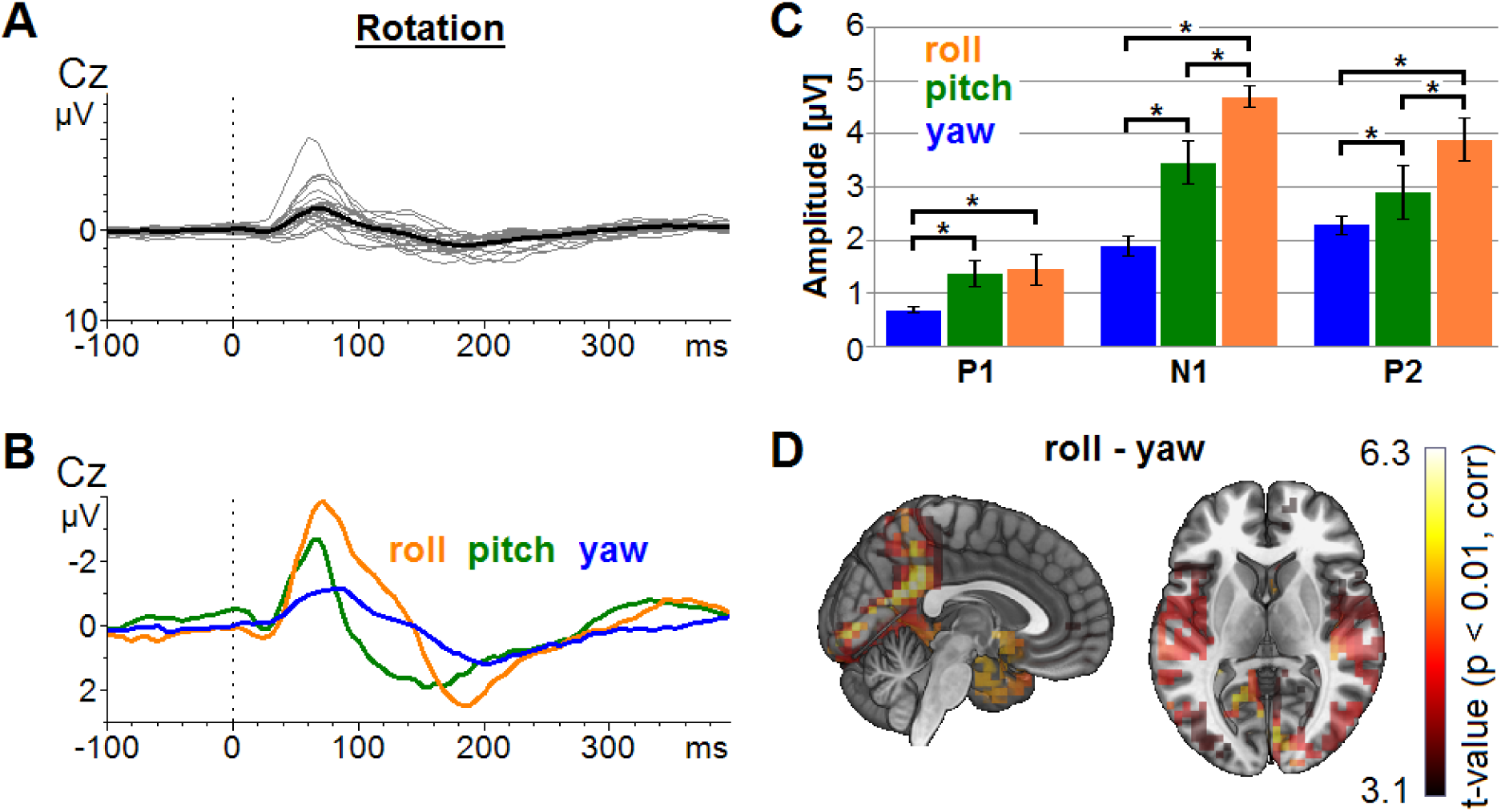
(A) Grand average ERPs (black line) for all rotations around the three main axes (yaw, pitch, roll). The grey lines represent the average ERPs of the single subjects for all three rotation axes. (B) Grand average ERPs elicited by roll, pitch, and yaw rotations. (C) The amplitudes of the VestEPs are modulated by the rotation axes. The whiskers represent the standard error of the mean. (D) A distributed source localization analysis was used to identify the neural generators of the ERPs displayed in panel B. According to this analysis, the amplitude differences between roll and yaw rotations are best explained by increased activity in cingulate gyrus elicited by roll movements.

Motion platforms consist of 6 actuators and the geometric arrangement determines the possible movements. Because arrangement is not point-symmetrical, the maximum displacement depends on the movement direction. For this reason, the maximum displacements and acceleration intensities for pitch and roll tilts differed significantly and since we could show in a previous study that the amplitudes of the VestEPs can be modulated by acceleration intensities [15] a systematic comparison of the tilt and translation amplitudes does not appear to be reasonable. The amplitudes evoked by roll movements were generally larger than those evoked by pitch tilt, which is in line with the previous increase by intensity findings (Fig. 3).

### ERPs – rotations

The grand average of the ERPs elicited by rotations also revealed three distinct components (Fig. 3). The P1 had an amplitude of 0.85 µV (SD = 0.59 µV) and peaked 29.7 ms (SD = 15.3 ms) after the acceleration maximum. The N1 component reached its maximum amplitude of -2.98 µV (SD = 1.98 µV) with a latency of 77.4 ms (SD = 12.3 ms). The third component had a mean amplitude of 2.41 µV (SD = 1.09 µV) and peaked with a latency of 169 ms (SD = 40.5 ms).

A comparison of the latencies and amplitudes across the three rotation axes revealed significant differences for the amplitudes for the P1 (F = 7.38, p = 0.001), the N1 (F = 12.1, p < 0.001), and the P2 (F = 6.53, p = 0.002) components. The post-hoc t-tests showed that the amplitudes for yaw rotations were significantly reduced for all three components compared to pitch (P1: p < 0.001; N1: p < 0.001; P2: p = 0.005) and roll (P1: p < 0.001; N1: p < 0.001; P2: p < 0.001) rotations. Also, the N1 and P2 amplitudes for pitch rotations were significantly smaller than those elicited by roll rotations (Fig. 3). No significant differences were found for the latencies.

### ERPs – component correlations

Here we investigated whether the amplitudes of the three components (P1, N1, P2) found in all subjects for all conditions (tilt, translation, rotation) correlated with each other. Thus, we calculated the Pearson correlation coefficients and corrected for multiple testing. Significant correlations between all components were found. The correlation between the amplitudes of the P1 and N1 was r = 0.307 (p = 0.013), between the N1 and P2 amplitudes r = 0.578 (p < 0.001), and the correlation between the P1 and P2 amplitudes was r = 0.283 (p = 0.022).

### Evoked Vestibular Beta-Band Response (EVBBR)

The analysis of the frequency space for all three stimulations consistently revealed an evoked vestibular beta band response elicited by accelerations (Fig. 4, bottom). As for the ERPs, the EVBBR power across directions was compared within the rotation condition. Significant differences for the EVBBR power (F = 6.82, p = 0.002) but not for the latencies (F = 2.11, p = 0.13) were found. The post-hoc t-tests revealed that the pitch (p = 0.002) and roll (p < 0.001) power is significantly larger than the yaw power, while the power did not differ between the rotations around the pitch and roll axes.

**Fig. 4:**
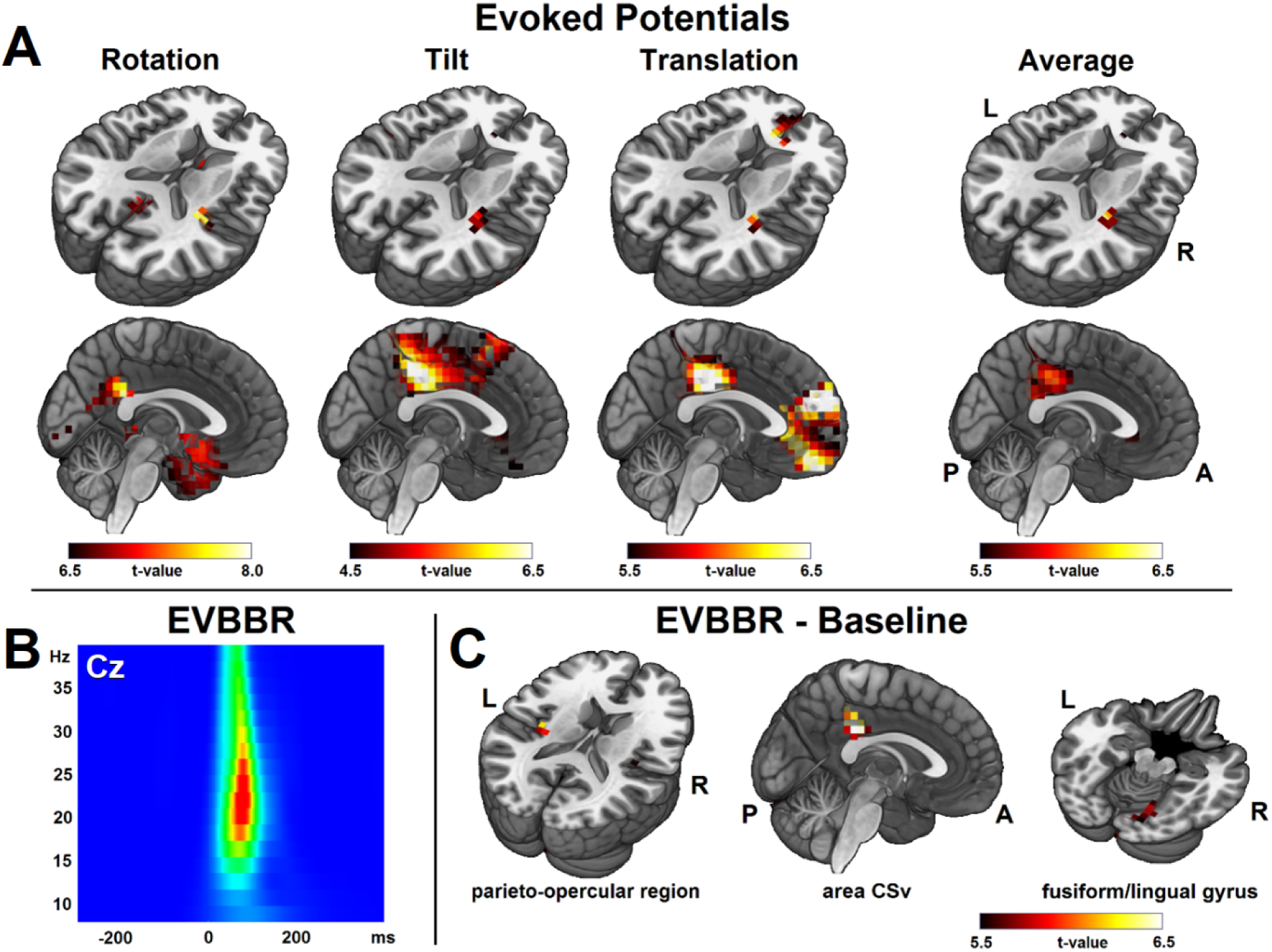
(A) Overlays of the distributed source localization by means of eLORETA for each stimulus profile onto an MNI template brain. The average activity was localized in areas well known as hubs of the cortical vestibular network, such as the parieto-opercular region and area CSv. (B) Visualization of the evoked vestibular beta-band response (EVBBR) visible as a transient power increase at electrode Cz. (C) Source level contrast indicating that the EVBBR is generated by increased beta activity in area CSv and the parieto-opercular region.

### ERP – EVBBR correlations

The mean EVBBR power across subjects was significantly correlated with the mean amplitudes of all three EEG components. The weakest correlation (r = 0.254, p = 0.041) was observed between the EVBBR and the P1 amplitude. The correlations between the N1 and P2 of r = 0.621 (p < 0.001) and r = 0.641 (p < 0.001) respectively were highly significant.

### Source localization – eLORETA

The distributed source localization of the evoked potentials in the time domain revealed similar patterns for all three stimulations. We found the right parieto-opercular region, area CSv, and Brodmann area 32 to be the most consistent sources (Fig. 4, top). Contrasting rotations and translations, area CSv was found to explain most of the difference with stronger activity during translations. Additionally, significantly more activity in Brodmann area 8 including the frontal eye field was observed during rotations. The activity in Brodmann area 8 as well as in the dorsal anterior cingulate gyrus (Brodmann area 32) was also significantly increased for rotations versus tilts. The contrast between translations and tilts revealed differences in several areas including the left- and right-sided parieto-opercular cortex, area CSv, Brodmann area 6, the medial frontal gyrus, and the inferior frontal gyrus. Differences between rotations and tilt/translations might be partially explained by the different visual input between both conditions (fixation versus eyes closed).

The distributed source localization of the EVBBR revealed a network containing area CSv, the bilateral parietal operculum, and the fusiform/lingual gyrus as the origin of the beta oscillations (Fig. 4, bottom).

### Direction-dependent differences

As stated before, the amplitudes of all three potentials were modulated by the rotation axes (yaw, pitch, roll). Particularly, increased amplitudes were observed for rotations in the roll plane. The results of the distributed source localization revealed that area CSv contributes the most to the signal increase (Fig. 3D). Analyzing the VestEPs evoked by opposing acceleration directions separately (fore vs. aft, right vs. left), no significant differences were found. Only The P1 amplitudes elicited by aft accelerations, were significantly reduced (p = 0.018, uncorr.) compared to fore accelerations (Fig. 5), indicating that the acceleration direction cannot be retrieved from the VestEPs.

**Fig. 5:**
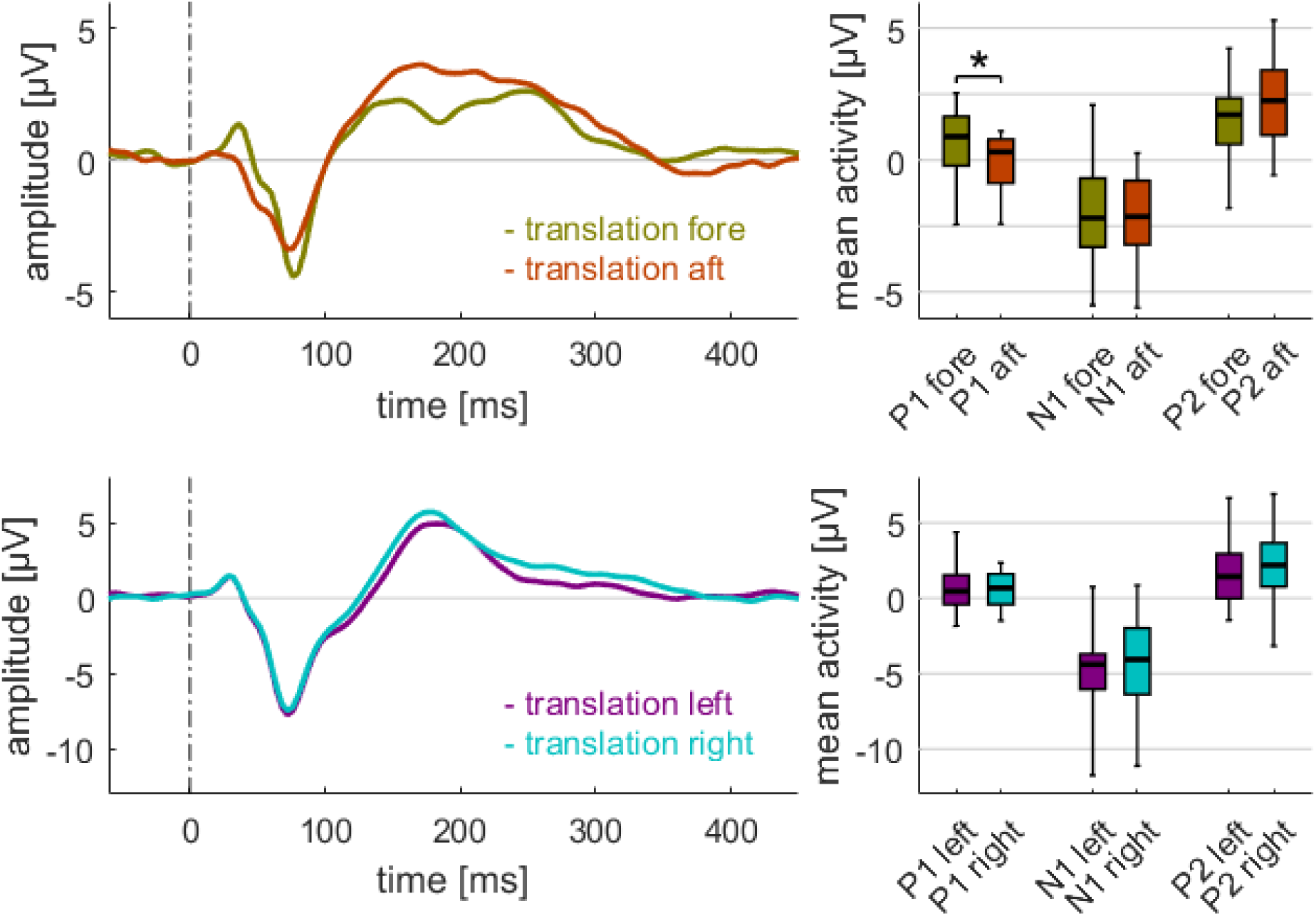
A sub-analysis for translations in the fore/aft (top row) and left/right direction (bottom row). Only the P1 amplitude with respect to the fore/aft direction differed when comparing the amplitudes within the motion axis. The time courses on the left represent the grand averages per condition.

### Combined Tilts and Translations

In addition to the isolated tilts, translations, and rotations, two conditions with an amalgamation of translation and tilt were performed for the pitch and roll plane. For one condition, the accelerations for combined tilt-translations added up, while in the second condition the movements for combined tilt-translations were chosen to cancel each other out in the optimal case and result in a net acceleration of zero. Reaching a net acceleration of zero at the position of the vestibular organs would require the knowledge of its exact position and orientation relative to the rotation axis of the platform. We did not individualize the motion profiles on a single subject level, which most likely resulted in residual non-zero accelerations in most subjects. The group comparisons between the “add” and the “cancel-out” condition revealed significantly reduced amplitudes (p < 0.001) for all three potentials in the cancel-out condition for the entire cortical vestibular network (Fig. 6). This plausibility check substantially illustrates and emphasizes the vestibular origin of the signal in contrast to potential somatosensory confounders of the stimulation.

**Fig. 6:**
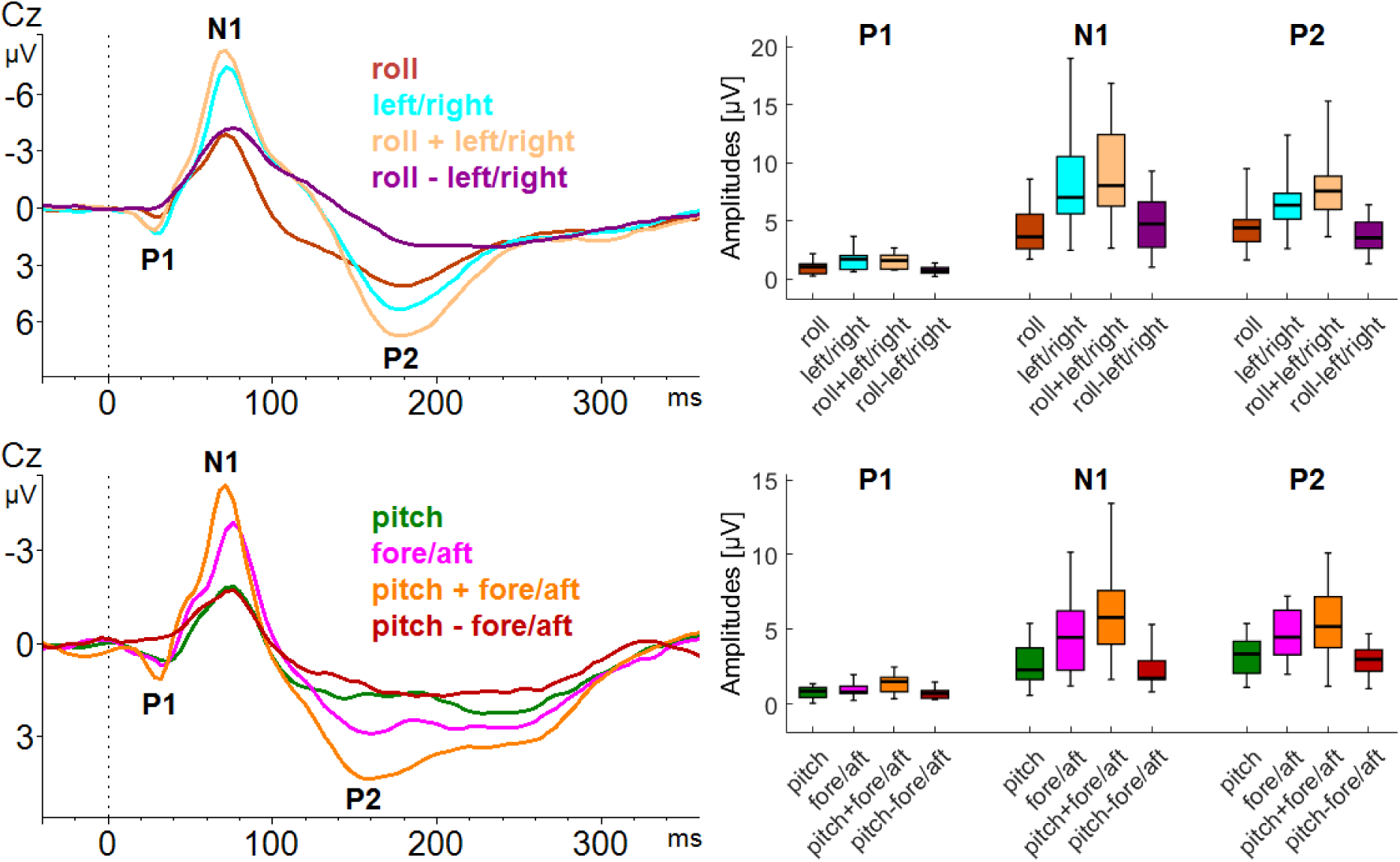
An analysis of isolated tilts and translations and combined tilt-translations in the roll (top) and pitch plane (bottom) revealed significantly reduced cortical amplitudes for conditions where the tilt and translation vectors partially cancelled each other out (roll – left/right and pitch – fore/aft) compared with the conditions in which both acceleration vectors added up (roll + left/right and pitch + fore/aft). The time courses on the left represent the grand averages per condition.

## Discussion

This study is the first EEG study to investigate the full spectrum of 3D head movements detectable by the vestibular organ in healthy humans. A recent finding of a distinctive evoked beta-band response to vestibular stimulation [15] could be replicated and consolidated for all accelerations applied in this study. In the current study, VestEPs were found to be influenced by the orientation of the rotation axis with respect to the head. We also could demonstrate that naturalistic rotations, translations, and tilts result in an overlapping cortical response pattern generated within a common network governed by the posterior perisylvian cortex and area CSv in the cingulate cortex.

### Amplitude modulation by acceleration direction

Significant differences in amplitudes were found for the three different rotation axes (yaw, pitch, roll). Based on the amplitude relationships, we assume that the difference might be linked to the occurrence of head rotation around these axes in daily life. In the absence of data, we speculate that the head rotations most frequently performed by humans are the rotations around the yaw axis, while rotations around the pitch and roll axes are less frequently performed. Unfortunately, no continuous tracking data on natural head motions in daily life is available [8]. One could argue that common movements are less important to cortical structures, while uncommon input needs a careful evaluation and interpretation by the cortical vestibular network and might therefore cause increased evoked potentials. It has been well studied in other contexts, such as oddball-experiments [17], that increased amplitudes can correspond to higher computational effort due to a more careful evaluation and integration of the incoming sensor information by the neo-cortex for novel stimuli.

The comparison of the fore and aft accelerations revealed only the P1 amplitude to significantly differ between the directions. This might be surprising at first glance, but it is important to keep in mind that the vestibular organ provides the brain with information on head accelerations. In daily life, accelerations typically occur only as acceleration-deceleration pairs with potentially differing acceleration profiles. In other words, every movement has a start and an end and the accelerations at the start and end have opposite signs.

An alternative explanation for alterations in the VestEP amplitudes can be derived from a study in which questionnaires, saliva cortisol and α-amylase were used to quantify mood states during passive motions [58]. The study reported that motion patterns were associated with specific mood stages. Particularly, yaw rotations were associated with feeling more comfortable while roll rotations with feeling less comfortable and pitch movements with a state of alertness. This provides an alternative explanatory framework for differences in amplitudes by an inter-play between the emotion and sensory systems.

### Source Localization and the role of area CSv

The results of the ERP source localization confirmed our previous findings [14–16] and demonstrated that EEG source localization can produce robust, meaningful, and reproducible results in the context of natural vestibular stimulation. In all conditions we found the parietal operculum and area CSv as the main hubs of the cortical vestibular network (Fig. 4). We argue that the differences between the conditions are more likely to reflect stimuli and task properties as well as natural priming rather than fundamental differences in the cortical processing of tilts, translations, or rotations.

One might wonder about the consistent finding of area CSv as one of the main hubs in the vestibular network in EEG studies, while the region was less frequently reported by other neuroimaging modalities in the past. Two different explanations are possible: Firstly, most fMRI and PET studies used block designs in which the vestibular stimulation was applied for multiple seconds, whereas we used extremely short stimuli in the EEG studies. It is possible that cortical vestibular processing is a two-stage “observe” and “assess” process. In this scenario, some areas might only contribute to vestibular processing during the initial assessment of a changed input stream. Given that area CSv is sensitive to acceleration intensity and acceleration direction, it might only be active for a limited time after the acceleration occurred. Similar two-step models have also been proposed in other fields of neuroscience, such as the implementation and maintenance model in emotion regulation [26,40]. The second explanation is related to the stimuli used in the various experiments. One of the most frequently used stimulation methods in fMRI experiments is galvanic vestibular stimulation. This method induces a feeling of dizziness with little or no perception of directionality. It is possible that area CSv is not involved in the processing of undirected vestibular input and is therefore less frequently reported by those fMRI and PET studies. Taking results of fMRI experiments on visual motion into account [51] it seems that CSv represents a multisensory area highly involved in the processing of ego motion consistent stimuli.

Activity in frontal structures, especially the anterior cingulate cortex as described here, has been previously reported by other EEG studies on the vestibular system [27,54,55]. These studies used a dipole fitting approach and described evoked potentials with latencies shorter than 30 ms. All studies cross-confirm the involvement of frontal structures probably during very early stages of cortical vestibular processing. Interestingly, frontal activity seems to be a less consistent finding in imaging studies. This might be due to excellent temporal resolution of the EEG which enables researchers to record hundreds of repetitions and allows short transient events to be detected.

### The Evoked Beta-Band Response (EVBBR)

Beside the description of VestEPs in the time domain, frequency analyses have also been performed in the past. One of the pioneering studies [2] reported strong changes in the delta and theta band during vestibular stimulation compared to a resting condition. The effects were most dominant over parietal and central structures. Additionally, in line with a recent study [18], a significant reduction in alpha power during vestibular stimulation was observed. No significant changes were reported for higher frequencies such as the beta- or gamma-band. It is important to note that both studies used prolonged accelerations different from the impulse-like accelerations used by us. Profiles without a clear motion onset introduce a temporal jitter, which particularly impacts the evoked response in high frequency bands. In our data, we found, motivated by earlier findings [15] an increased EVBBR for all three types of stimuli (rotations, translations, tilts) and the EVBBR power was closely linked to the N1 and P2 amplitudes by means of a correlation analysis. The EVBBR showed a stronger correlation to the N1 and P2 potentials than the ERP components show to each other. This can lead to the interpretation that the beta oscillation generating neural process feeds into or even drives the ERP generating processing steps. The distributed source localization of the EVBBR suggests that beta oscillations might be involved in long-range interaction between the left and right parieto-opercular regions and area CSv. The beta-band activity in the right fusiform/lingual gyrus is more challenging to interpret. The activity is at the edge of the electrical head space model (cerebral cortex) and could very well reflect mislocalized vestibular stimulation response activity originating in the vestibulo-cerebellum (anterior vermis).

### Rotations versus Translations

A quantitative comparison of the VestEPs across the stimulation conditions in the context of this study would be problematic. The reason for this is that the exact accelerations sensed by the different parts of the vestibular organs are, due to geometric uncertainties in our experimental setup, hard to estimate and a comparison between the conditions might mainly reflect unequal stimulus intensities. Therefore, only a qualitative comparison of the time courses was performed. Here, the time courses elicited by translations, tilts, and rotations as well as the estimated sources for all three conditions showed very similar patterns (Fig. 3, 4). It is very likely that vestibular input causes more specific direction and movement responses in infratentorial regions of the central vestibular network.

The comparisons between the “add” and the “cancel-out” condition for the combined tilt and translation movements showed reduced amplitudes for all potentials in the “cancel-out” condition (Fig. 6). This difference might point to the vestibular origin of the observed ERPs in contrast to potential somatosensory confounders of the stimulation. It is important to keep in mind that cancelation only refers to the acceleration detected by the otolith organs. Accelerations detected by the semicircular canals and the forces acting on any other part of the body do not add to zero. The ERPs visualized in figure 6 show that for both movement planes the combined tilt + translation results in the largest amplitudes, while the tilt – translation amplitudes are of the same magnitude as pure tilt profiles. Since the forces acting on the body during the tilt +translation movement are larger compared to the once occurring during pure tilts, it is reasonable to argue that the contribution of somatosensory processing to the VestEPs is limited.

### Limitations

Eye movements were not recorded in either of the two experiments and hence the possibility that despite a careful state-of-the-art artifact correction in preprocessing of the data, some acceleration-correlated eye movements may still have contaminated the VestEPs cannot be completely ruled out. However especially the impact of ocular counter rolling should be negligible since the artifact causing the dipole does not change position relative to the sensors in such situations.

The different eye states, open eyes in experiment 1 and eyes closed in experiment 2, might hamper a between-experiment comparison. Though, earlier experiments [15] systematically investigated the impact of such differences and did not find any significant impact of visual input (fixation/eyes open in dark/blindfolded) on VestEPs.

We cannot completely rule out alternative explanations for the correlations between the VestEPs and the EVBBR such as impedance or a spurious correlation due to an overall power increase.

## Acknowledgments

Part of this work was prepared in the context of Marie Woller’s dissertation at the Faculty of Medicine, Ludwig-Maximilians-University Munich. This work was supported by the German Federal Ministry of Education and Research (German Center for Vertigo and Balance Disorders - IFB^LMU^, Grant code 01EO140) and the German Foundation for Neurology (Deutsche Stiftung Neurologie). The authors would like to thank Katie Göttlinger for copy-editing the manuscript.

## Data Availability

The data sets generated and/or analyzed during the current study are available from the corresponding author on reasonable request.

